# ABLs and TMKs are co-receptors for extracellular auxin

**DOI:** 10.1101/2022.11.28.518138

**Authors:** Yongqiang Yu, Wenxin Tang, Wenwei Lin, Wei Li, Xiang Zhou, Ying Li, Rong Chen, Rui Zheng, Guochen Qin, Wenhan Cao, Patricio Perez, Rongfeng Huang, Jun Ma, Juncheng Lin, Liwen Jiang, Tongda Xu, Zhenbiao Yang

## Abstract

Extracellular perception of auxin, an essential phytohormone in plants, has been debated for decades. Auxin binding protein 1 (ABP1) physically interacts with quintessential transmembrane kinases (TMKs) and was proposed to act as an extracellular auxin receptor, but its role was disputed because *abp1* knockout mutants lack obvious morphological phenotypes. Here we identified two new auxin-binding proteins, ABL1 and ABL2, that are localized to the apoplast and directly interact with the extracellular domain of TMKs in an auxin-dependent manner. Furthermore, functionally redundant ABL1 and ABL2 genetically interact with TMKs and exhibit functions that are overlapping with those of ABP1 as well as independent of ABP1. Importantly, the extracellular domain of TMK1 itself binds auxin and synergizes with either ABP1 or ABL1 in auxin binding. Thus, our findings discovered new auxin receptors ABL1 and ABL2 having functions overlapping with but distinct from ABP1 and acting together with TMKs as co-receptors for extracellular auxin.

## INTRODUCTION

Auxin regulates nearly every aspect of plant growth, development, and responses to the environment, and has been shown to control transcription-independent rapid responses occurring in seconds as well as slow transcriptional responses ^1–7^. The latter requires the intracellular receptors TRANSPORT INHIBITOR RESPONSE 1 (TIR1)/AUXIN-SIGNALING F-BOX proteins (AFBs) that promote the degradation of their co-receptors AUX/IAA transcriptional repressors and act as adenylate cyclases ^8–13^. Increasing studies show that many auxin-induced effects such as plasma-membrane hyperpolarization, cytosolic Ca^2+^-transients, activation of ROP GTPases, RAF-like kinase-dependent cytoplasmic streaming, and protoplast swelling, are too rapid to depend on transcriptional regulation ^5,7,14–21^. Most of these rapid auxin responses are evidently regulated by a class of auxin receptors distinct from the nuclear TIR1/AFB receptors and presumably localized to the cell surface ^7,14,22,23^.

A key hallmark of auxin action lies in its dynamic flux across the plasma membrane and the cell wall of adjacent cells and also its long-distance flow across tissues and organs, which relies on the intercellular flux ^24–29^. Thus, it is imperative that apoplastic auxin levels are tightly monitored and controlled. Furthermore, cell surface auxin signaling is needed to rapidly regulate cytoplasmic responses when extracellular auxin levels change. Indeed, the transmembrane kinase (TMK) family receptor-like kinases are the essential plasma membrane-localized auxin signaling components required for many auxin responses including rapid responses such as cell wall acidifications, ROP GTPase activation, and proteins phosphorylation ^3,14,15,30,31^. Auxin activates TMK phosphorylation, and the activated TMK kinase domain directly phosphorylates a series of effectors such as IAA32/34, AHA, MKK4/5-MPK3/6, TAA1, and ABI1/2 to regulate multiple developmental processes ^14,32–35^. However, the molecular mechanism for the perception of auxin signals that activate TMK-dependent auxin responses remains enigmatic. Auxin-binding protein 1 (ABP1) forms a complex with the extracellular domain of TMK1 in an auxin-dependent manner, suggesting that it may sense auxin to activate TMK-dependent auxin responses ^15^. However, the role of ABP1 in auxin perception/signaling was debated, because knocking out the single-copy *ABP1* gene did not produce visible phenotypes in *Arabidopsis* ^36^. In contrast, knocking out all four functionally redundant TMKs, as in the *tmk1;tmk2;tmk3;tmk4* mutant, induces a wide range of phenotypes including embryo and seedling lethality in *Arabidopsis* ^15,37^. A recent study shows that both *abp1* and *tmk1* null mutants exhibit similar defects in auxin-induced rapid global phospho-responses and canalization in *Arabidopsis* and that auxin binding to ABP1 is required for these auxin responses, corroborating the function of ABP1 in TMK1-dependent auxin perception/signaling ^3,23^. Yet this study fails to explain why many TMK-dependent developmental processes and auxin responses are unaffected in the *abp1* null mutants. Importantly whether and how the quintessential TMKs perceive extracellular auxin remains unresolved.

Given the lack of obvious phenotypes in *abp1* knockout mutants and the predominant localization of ABP1 to ER ^3,36,38^, we speculated the existence of other extracellular auxin-binding proteins that might act together with TMKs to perceive extracellular auxin. To search for such proteins, we conducted an immunoprecipitation-mass spectrometry analysis from a TMK1-Flag *Arabidopsis* line and identified two highly related germin-like proteins that contain an auxin-binding box-like motif similar to A-box found in ABP1. Here we demonstrate that these ABP1-like (ABL) proteins and TMKs act as co-receptors for sensing extracellular auxin.

## RESULTS

### The auxin-binding pocket mutant protein ABP1-5 suppresses auxin responses

To assess the existence of other auxin binding proteins acting with TMKs, we first investigated the effect of *ABP1-5*, harboring H94Y mutation in its auxin binding pocket, which shows compromised auxin-induced interaction with TMK1 ^15^. Expressing *ABP1-5 in vivo* is anticipated to cause a dominant negative effect on other auxin-binding proteins interacting with TMKs. We generated *gABP1-5* (*pABP1::ABP1-5)* lines in the *abp1* (*abp1-TD1*) knockout mutant ^36^ background and analyzed their phenotypes in comparison with *gABP1;abp1* (*pABP1::ABP1;abp1-TD1)*. ABP1 and ABP1-5 protein levels in these lines were comparable (Figure S1A). *gABP1-5;abp1* showed a series of growth and developmental phenotypes and was compromised in auxin responses, which were not observed in either *gABP1;abp1,* or wild type. As in the *tmk1;tmk2;tmk3;tmk4* mutants ^15^, the auxin-induced lobing of pavement cells in cotyledons was compromised in *gABP1-5;abp1* but not in *gABP1;abp1* (Figures 1A-C). We previously showed that auxin promotes lobe and indentation formation by activating ROP2 and ROP6, respectively ^2^. Consistently, the auxin-promoted activation of both ROP2 and ROP6 was partially abolished in *gABP1-5;abp1* but not in *gABP1;abp1* (Figures 1D-F). As in the *tmk1;tmk4* mutant ^14,15^, auxin-promotion of hypocotyl elongation and cotyledon epinasty was attenuated in *gABP1-5;abp1* but not in *gABP1;abp1* (Figures 1G-I). Furthermore, PIN1 protein was depolarized and internalized in the root cells of the *gABP1-5;abp1* transgenic lines (Figures S1B and S1C), agreeing with reduced root length and gravitropic responses (Figures S1D-G) as in *tmk1;tmk4* mutants ^34^. *gABP1-5;abp1* lines displayed altered cotyledon developmental patterns including single and triple cotyledons, as in *pin1* ^39^ and *tmk1;tmk2;tmk3;tmk4* ^15^ (Figure S1H). *gABP1-5;abp1* rosette leaves also exhibited abnormal shapes as found in the *pin1* mutant (Figure S1I). None of the phenotypes described above were observed in the *abp1* mutants or the *gABP1;abp1* lines (Figures S1D-I). Altogether, our results show that the accumulation of the ABP1-5 protein impacts TMK-modulated auxin responses likely by causing a dominant negative effect on other auxin-binding proteins that functionally compensate for the loss of ABP1.

**Figure 1.**
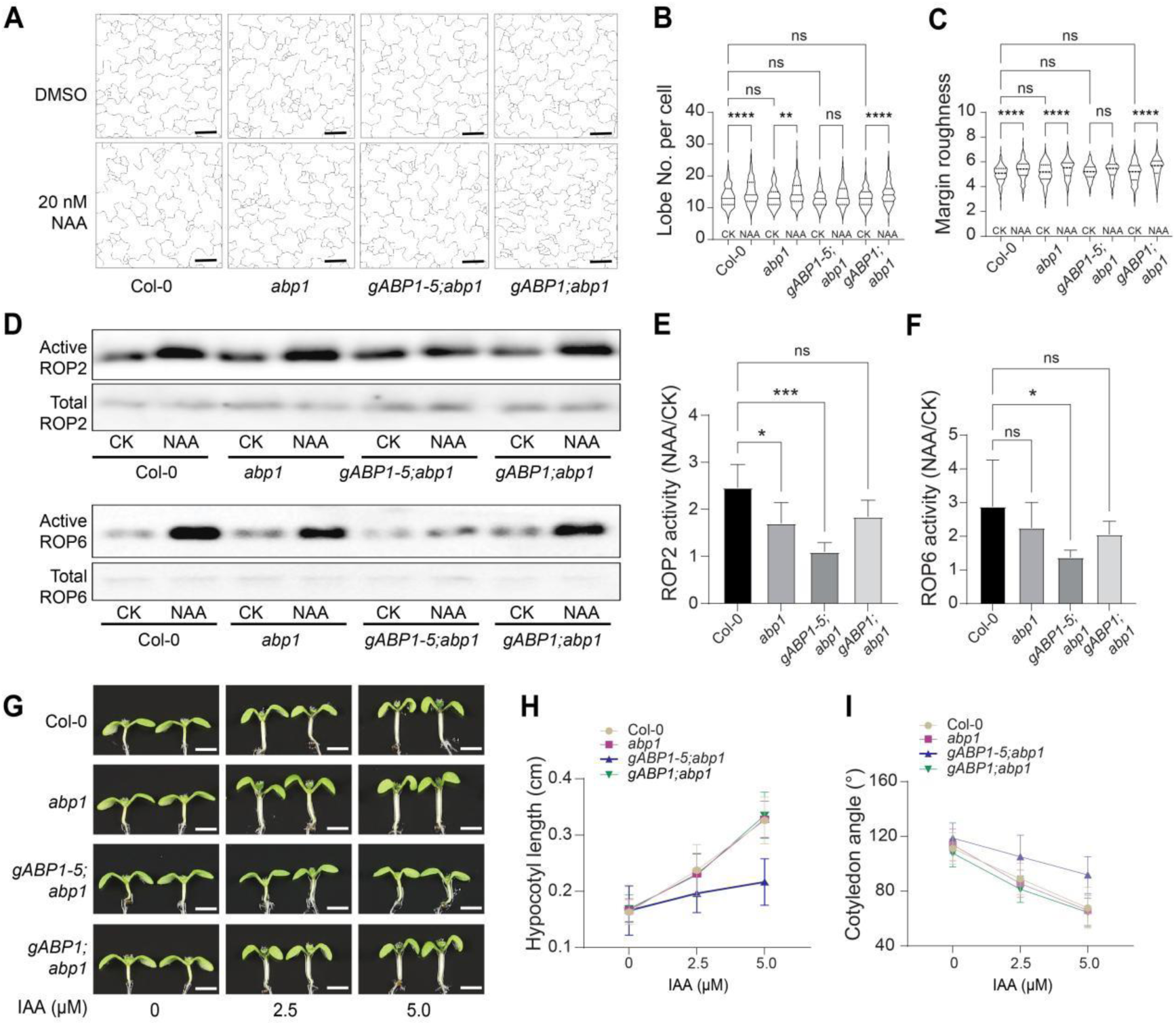
ABP1-5 suppresses auxin responses in *Arabidopsis*. Representative auxin responses in Col-0, *abp1*, *gABP1-5;abp1* and *gABP1;abp1* lines are presented here.(A) The phenotype of epidermal pavement cells (PC) in *Arabidopsis* cotyledons of Col-0, *abp1*, *gABP1-5;abp1* and *gABP1;abp1* with mock (DMSO) and NAA treatment. Scale bar, 50 µm. (B-C) Quantitative analysis of PC interdigitation is depicted by the number of lobes per cell (B) and margin roughness (C). Cotyledons from different lines were treated with DMSO and NAA as described in (A). n>147 independent cells for each treatment. ns denotes not significant; \*\**p < 0.01*;\*\*\*\**p < 0.0001*; one-way ANOVA.(D-F) ROP2 and ROP6 activity assays using the protoplasts of Col-0, *abp1*, *gABP1-5;abp1* and *gABP1;abp1* treated with 50 nM NAA for 10 mins. Active ROP2 (E) and ROP6 (F) levels (the amount of GTP-bound ROP2 or RO6 divided by the amount of total ROP2 or ROP6) were measured. Auxin-induced activity (NAA) relative to mock control (CK, designated as ‘ 1’’) is shown. Data are mean ± SD (n=4 independent experiments). ns denotes significant; \**p < 0.05*; \*\*\**p < 0.001*; one-way ANOVA.(H-I) Auxin-promotion of hypocotyl elongation and cotyledon bending in Col-0, *abp1*, *gABP1-5;abp1* and *gABP1;abp1*. 5-day-old seedlings were treated with 0, 2.5 and 5 μM IAA, respectively, and hypocotyl length (H) and cotyledon angles (I) were measured and quantified. Scale bar, 2 mm. Data are mean ± SD (n>28 independent seedlings).

### ABP1-like proteins (ABLs) form auxin sensing complex with TMKs in the apoplast

Because we failed to identify clear *ABP1*-homologous genes in the *Arabidopsis* genome by amino acid sequence homology search, we explored potential new auxin binding proteins that compensate for ABP1 in TMK-mediated auxin signaling by immunoprecipitating the TMK complex from the *pTMK1::TMK1-Flag* transgenic line. The immunoprecipitants contained a member of the cupin protein family where ABP1 belongs, which we termed <u>AB</u>P1-like protein 1 (ABL1) (Supplemental Table 2). By sequence homology search, we identified ABL2 from the *Arabidopsis* genome. *ABL1* (AT1G72610) and *ABL2* (AT5G20630) encode polypeptides with 208 and 211 residues respectively, sharing high amino acid sequence similarity with each other (64.42% identity). ABL1 and ABL2 exhibit low sequence similarity to ABP1 (26.26% and 18.44% identity, respectively), but contain a conserved auxin binding pocket with key amino acids predicted to be involved in auxin binding ^40,41^ (Figure S2A). AlphaFold-based protein structure simulations predicted a three-dimensional structure for ABL1, which shares high similarity with that for ABP1, especially in the auxin binding pocket (Figure S2B). Structure-based blind docking of ABL1 and ABL2 protein predicted that this pocket binds NAA, suggesting a potential role for ABL1 and ABL2 as new auxin binding proteins (Figure S2B).

To assess whether both ABL1 and ABL2 form auxin sensing complexes with TMKs, we generated anti-ABL1 and anti-ABL2 antibodies that specifically detected ABL1 and ABL2 proteins, respectively (Figure S2C). ABL1 and ABL2 are predicted to be secreted as they contain signal peptides and lack other targeting signals. We first analyzed the subcellular localization of ABL1 in *Arabidopsis* by both immuno-fluorescence staining and immuno-gold labeling. Both assays showed that ABL1 is almost exclusively localized to the cell surface (Figures 2A and 2B and S2D-G), contrasting to ABP1, which contains an ER retention signal and is primarily localized to ER with a small fraction (10-15%) found in the apoplast ^3,38^. Immunogold labeling further determined that ABL1 is localized to the apoplast (Figure 2A).

**Figure 2.**
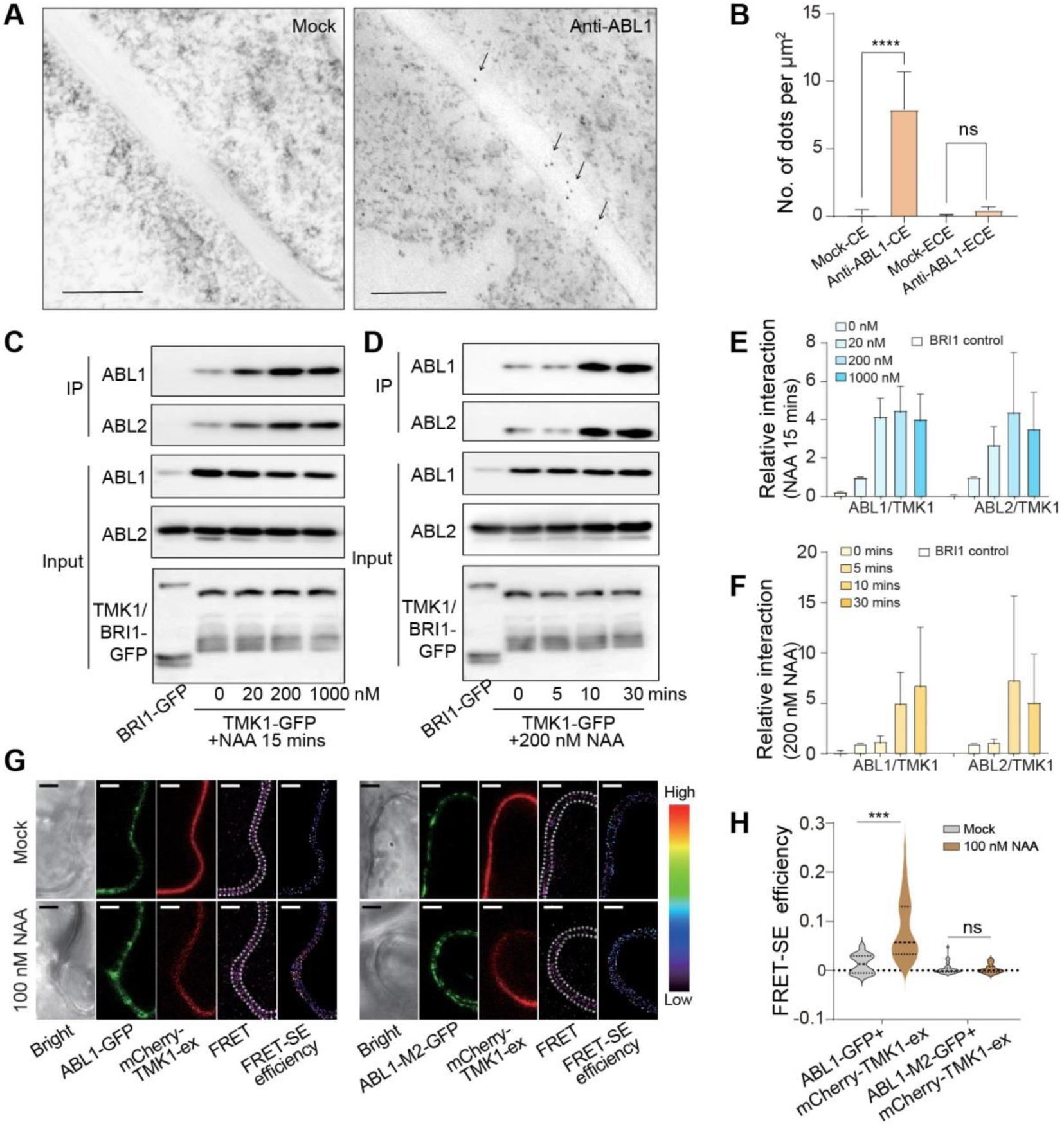
ABL1/ABL2-TMKs auxin sensing complexes on the cell surface. (A-B) The ABL1 protein localization in *Arabidopsis* young leaves was detected by the immune colloidal gold technique (A) and quantification (B). CE, cell wall, and extracellular region. ECE, regions except for cell wall and extracellular region. Scale bar, 500 nm. Data are mean ± SD (n>10 regions). ns denotes not significant; \*\*\*\**p < 0.0001*; one-way ANOVA.(C-D) The interaction between ABL1/ABL2 and TMK1 by co-immunoprecipitation from *Arabidopsis* seedlings expressing *pTMK1::TMK1-GFP* treated with different concentrations of NAA for 15 mins or treated with 200 nM NAA for different time periods. Proteins from TMK1-GFP and BRI1-GFP (as a negative control) seedlings were immunoprecipitated using GFP-Trap and tested by western blotting analysis using anti-ABL1 and anti-ABL2 antibodies (top two panels), respectively. Input amounts for ABL1, ABL2, TMK1-GFP and BRI1-GFP are shown (in middle and bottom panels).(E) Quantitative analysis of ABL1, ABL2 and TMK1 association before and after treatments with indicated concentrations of NAA for 15 mins in (C). Data are mean ± SD (n=3 independent experiments).(F) Quantitative analysis of the interactions between ABL1/ABL2 and TMK1 before and after treatments with 200 nM NAA for different time periods in (D). Data are mean ± SD (n=3 independent experiments).(G-H) FRET analysis between ABL1-GFP and mCherry-TMK1-ex in tobacco leaves. The representative heatmap of sensitized emission efficiencies of FRET between mCherry-TMK1-ex and ABL1-GFP or ABL1-M2-GFP (G). Images were obtained from the cell boundary region (dotted lines). Scale bar, 10 μm. Quantitative analysis of changes in the FRET-SE efficiency after 100 nM NAA treatment for 5 mins (H). n=45 cells for each treatment. ns denotes not significant; \*\*\**p < 0.001*; two-sided Student’s *t*-test.

The localization of ABL1 to the apoplast provides the spatial possibility to form a complex with TMK on the cell surface. Co-immunoprecipitation assays showed that both ABL1 and ABL2 form protein complexes with TMK1 in plant cells, respectively. Both ABL proteins were immunoprecipitated by anti-GFP antibodies from *TMK1-GFP* (*pTMK1::TMK1-GFP*) seedlings but not with BRI1 from *BRI1-GFP* seedlings (Figures 2C-D). The interaction between TMK1 with ABL1/ABL2 was enhanced within a few minutes by auxin treatments in a concentration-dependent manner with an optimal NAA concentration of 200 nM (Figures 2C-F), agreeing with its optimal concentration that activates ROP2 and ROP6 ^2,15^.

Fluorescence resonance energy transfer (FRET) analysis was used to investigate whether ABL1 directly interacts with the extracellular domain of TMK1 *in vivo*. We transiently expressed ABL1-GFP and mCherry-TMK1-ex (extracellular domain of TMK1) proteins in tobacco leaves. FRET signal was detected between ABL1-GFP and mCherry-TMK1-ex, and was strongly promoted by treatments with 100 nM NAA within a few minutes (Figures 2G-H). This auxin-induced interaction was completely abolished when two histidine residues (H100 and H102) in the putative auxin binding pocket were mutated to Ala (A) on the ABL1 protein (ABL1-M2) (Figures 2G and 2H). Consistent with the prediction that these two residues are essential for auxin binding, we found that these mutations indeed abolish ABL1’s ability to bind auxin (see below). Taken together our results indicate that secreted ABL1 and ABL2 directly interact with the extracellular domain of TMK1 and that this interaction is promoted by auxin. Hence, we hypothesize that the ABLs and TMKs form auxin receptor complexes on the cell surface to perceive extracellular auxin.

### Functionally redundant ABL1 and ABL2 show distinct and overlapping functions with ABP1

Because both ABLs and ABP1 form complexes with TMK1 in an auxin-dependent manner (Figure 2) ^15^, we asked whether ABLs and ABP1 are functionally redundant or distinct. We obtained both *abl1* (*abl1-1*) and *abl2* (*abl2-1*) single knockout mutants in the *A. thaliana* Col-0 background generated by either T-DNA insertion or CRISPR-based method (Figures S3A and S3B), but neither exhibited obvious defects in growth and development (Figure 3A). However, *abl1/2* (*abl1-1;abl2-1*) double mutant showed growth and developmental defects including reduced seedling size, curling leaves, and altered shapes of pavement cells (Figure 3A-D), suggesting functional redundancy between ABL1 and ABL2.

**Figure 3.**
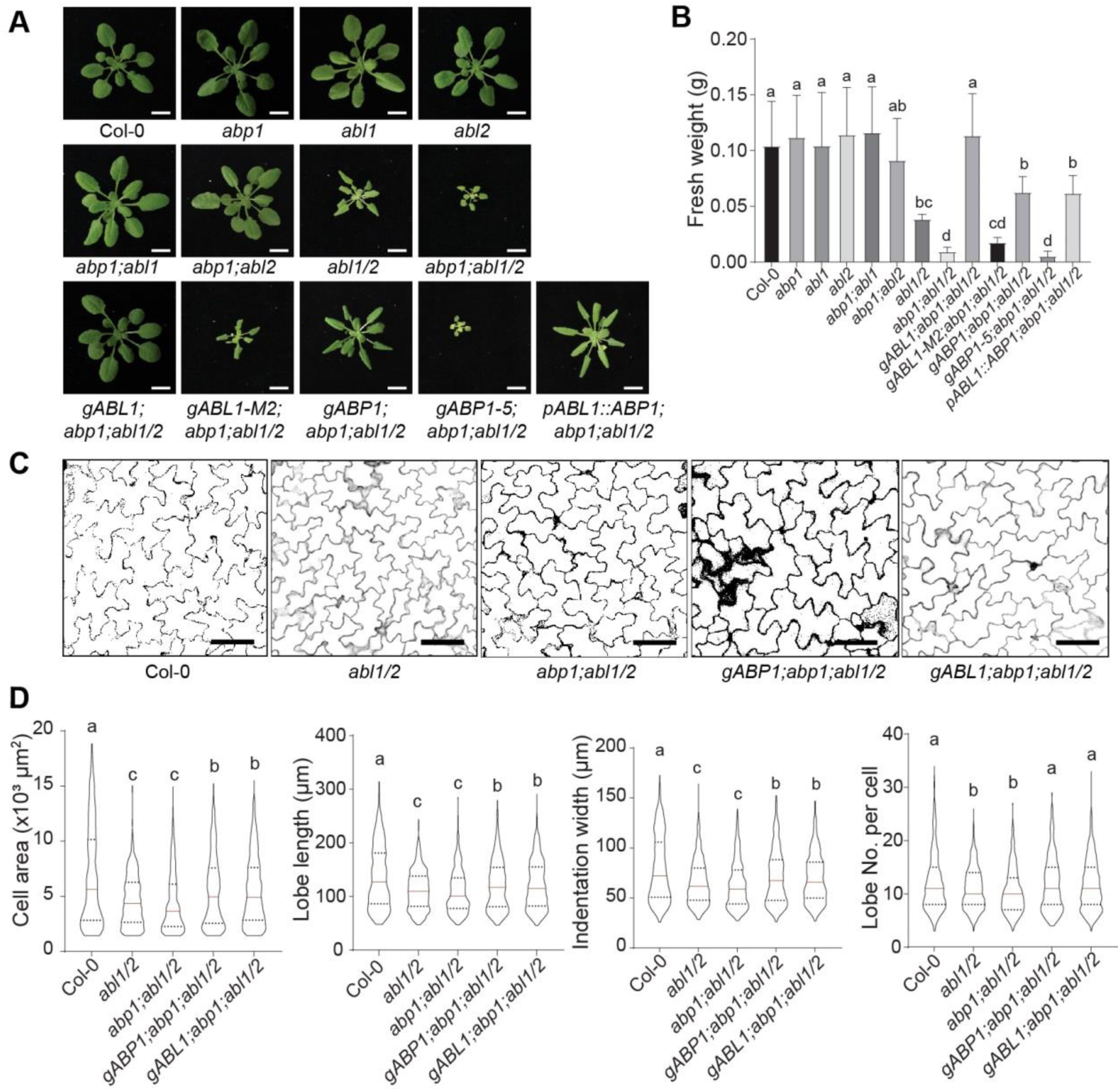
ABLs and ABP1 exhibit overlapping and distinct functions in plant development. (A) The morphology of 4-week-old soil-grown seedlings from Col-0, *abp1*, *abl1*, *abl*2, *abp1;abl1*, *abp1;ab12*, *abl1/2*, and *abp1;abl1/2*. The *abp1;abl1/2* phenotype was rescued by *gABL1*, *gABP1,* or *pABL1::ABP1*, but not by *gABP1-5* and *gABL1-M2*. Scale bar, 1 cm. (B) Quantitative analysis of fresh weights of seedlings as described in (A). Data are mean ± SD (>16 independent seedlings). Different letters indicate values with statistically significant differences (*p* < *0.05*; Tukey HSD).(C) Phenotype of pavement cells in 3-week-old the fifth pair true leaves of Col-0, *abl1/2, abp1;abl1/2*, *gABP1;abp1;abl1/2* and *gABL1;abp1;abl1/2*. Scale bar, 100 µm. (D) Quantitative analysis of PC interdigitation. Pavement cell area, length, indentation width, and the number of lobes per cell were analyzed. Data are mean ± SD (n>332 independent cells). Different letters indicate values with statistically significant differences (*p* < *0.05*; Tukey HSD).

To assess whether ABP1 is functionally redundant with ABL1 and/or ABL2, we generated *abp1-TD1;abl1-1* (*abp1;abl1*), *abp1-TD1;abl2-1* (*abp1;abl2*) double mutants and a *abp1-TD1;abl1-1;abl2-1* (*abp1;abl1/2*) triple mutant. The *abp1;abl1* mutant did not exhibit an obvious phenotype, while the *abp1;abl2* double mutant showed a slight reduction in seedling growth, much weaker than that in the *abl1;abl2* double mutant (Figures 3A and 3B). The *abp1;abl1/2* triple mutant showed greatly reduced plant sizes and curling leaves, much more severe growth and developmental defect than either *abl1;abl2,* or *abp1;abl1*, or *abp1;abl2* (Figures 3A and 3B), suggesting that ABP1 contributes to the ABL1/2-mediated processes. Growth defects such as small plant sizes in the triple mutants were nearly fully complemented by *pABL1::ABL1* but partially complemented by *pABP1::ABP1* or *pABL1::ABP1* (Figures 3A and 3B). Furthermore, some phenotypes such as leaf curling were complemented neither by *gABP1* nor by *pABL1::ABP1* but were fully complemented by *gABL1* (Figure 3A). These results show that the redundant ABL1 and ABL2 functionally overlap with ABP1 but also have ABP1-independent physiological functions. Importantly, ABL1 and ABL2 are the major contributors to the overlapping functions compared to ABP1, consistent with the differences in their subcellular distribution, and expression patterns (Figures S3C and S3D).

The *abp1-TD1;abl1-1;abl2-1* (*abp1;abl1/2*) triple mutant exhibited defects in pavement cell morphogenesis as well, which was partially complemented by *pABL1::ABP1* (Figures 3C and 3D). This triple mutant also displayed PIN1 localization defects as in *gABP1-5;abp1* (Figures S3E and S3F) or *tmk1;tmk2;tmk3;tmk4* mutants ^15^. Consistent with the defects of pavement cell morphogenesis and PIN localization, the activation of both ROP2 and ROP6 by auxin was partially abolished in the *abp1;abl1/2* mutant, which was partially restored by *gABP1* (Figures 4A and 4B). Furthermore, the *abp1;abl1/2* mutants showed insensitivity to the auxin promotion of hypocotyl elongation like both *gABP1-5;abp1* and *tmk1;tmk4*, and this *abp1;abl1/2* defect was rescued by *gABP1* (Figures 4C and 4D). These results confirm the function of ABP1 in the auxin-regulated pavement cell morphogenesis as previously reported ^2^, and further support its overlapping function with ABL1 and ABL2 in both plant development and auxin signaling.

**Figure 4.**
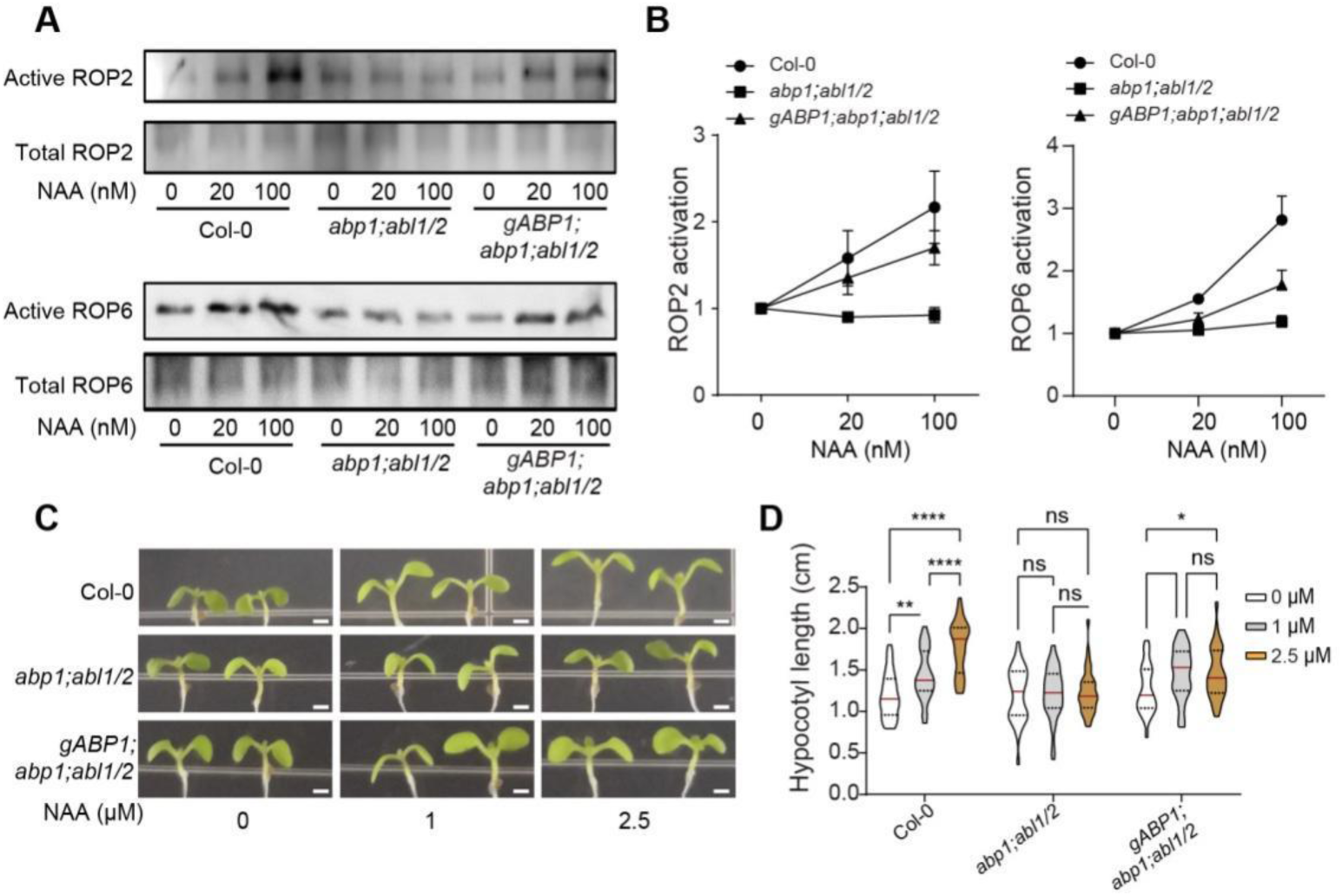
ABP1 and ABLs exhibit an overlapping function in the regulation of auxin responses. (A) ROP2 (top panel) and ROP6 (bottom panel) activation by auxin in Col-0, *abp1;abl1/2*, *abp1;abl1/2* and *ABP1* complementation line *gABP1;abp1;abl1/2* as described in Figure 1D. (B) The relative ROP2 and ROP6 activity levels in (A) were quantified by ImageJ software by measuring the intensity of the bands. The activity is the amount of GTP-bound ROP (top) divided by total ROP (bottom). Data are mean ± SE (n = 6 independent experiments).(C-D) Auxin-induced hypocotyl elongation in Col-0, *abp1;abl1/2* and *ABP1* complementation line *gABP1;abp1;abl1/2* (C). The hypocotyl length was quantified in (D). *n* = 30 independent seedlings per line, \**p < 0.05*; \*\**P* ≤ *0.01*, \*\*\*\**P* ≤ *0.0001*; ns denotes not significant, two-way ANOVA.

### Both ABL1 and ABP1 functionally interact with TMKs

Given the auxin-dependent physical interactions between ABP1/ABLs and TMKs, we were interested in their functional relationship in the regulation of auxin responses. We analyzed this relationship by respectively crossing *abp1, abl1,* and *abp1;abl1* with *tmk1-1-/+;tmk4-1 (tmk1-/+;tmk4)* heterozygous double mutant, which exhibited mild growth defects (Figures 5A and 5B). Neither *abp1* nor *abl1* enhanced the growth defects in *tmk1-/+;tmk4*, but the *abp1;abl1;tmk1-/+;tmk4* quadruple mutants showed a series of severe growth and developmental defects, much stronger than *abp1;abl1* or *tmk1-/+;tmk4,* including young seedling size (Figures 5A and 5B), plant architecture (Figure S4A), and inflorescence architecture (Figure S4B). All these developmental defects were restored by either ABP1 or ABL1 (Figures 5A, 5B and S4). More importantly, auxin promotion of hypocotyl elongation (Figures 5C and 5D), pavement cell morphogenesis (Figures 5E, 5F and S4C), and ROP activation (Figures 5G and 5H) were abolished in this quadruple mutant, and all these defects were rescued either by ABP1 or ABL1 (Figures 5C-H). Finally, we analyzed PIN1 localization in these mutants by immuno-staining, and found that PIN1 was depolarized in the *abp1;abl1;tmk1-/+;tmk4* mutant (Figures S4D and S4E). Moreover, *abp1;abl1;tmk1-/+;tmk4* showed reduced primary root length compared with the double or triple mutants (Figures S4F and S4G). These data show that functionally overlapping ABP1 and ABL1 genetically interact with TMK1 and TMK4 in regulating both plant development and auxin responses. Together with the auxin-dependent physical interactions between ABLs/ABP1 and TMK1, our findings conclusively demonstrate that ABLs/ABP1 and TMKs act together on the cell surface in the regulation of cytoplasmic auxin responses.

**Figure 5.**
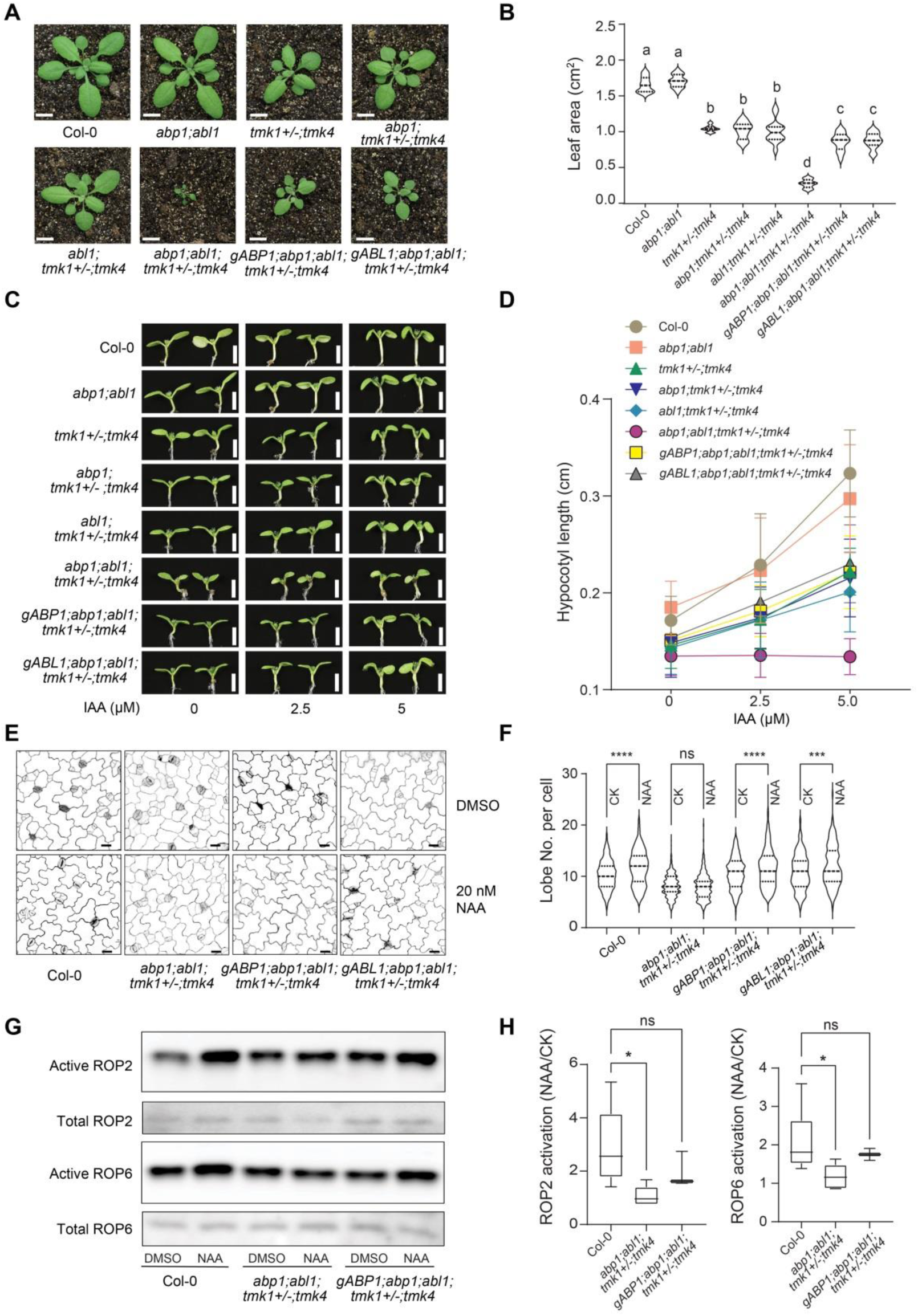
ABL1 and ABP1 function through their interaction with TMKs. (A) The seedling morphology of 4-week-old soil-grown Col-0, *abp1;abl1*, *tmk1+/-;tmk4*, *abp1;tmk1+/-;tmk4*, *abl1;tmk1+/-;tmk4*, *abp1;abl1;tmk1+/;-tmk4*, *gABP1;abp1;abl1;tmk1+/-;tmk4* and *gABL1;abp1;abl1;tmk1+/-;tmk4* lines. Scale bar, 1 cm. (B) Quantitative analysis of leaf area from the fifth true leaf of seedlings shown in (A). Data are mean ± SD (n=14 leaves from independent seedlings). Different letters indicate values with statistically significant differences (*p < 0.05*; Tukey HSD).(C) Auxin-induced hypocotyl elongation in Col-0, *abp1;abl1*, *tmk1+/-;tmk4*, *abp1;tmk1+/-;tmk4*, *abl1;tmk1+/-;tmk4*, *abp1;abl1;tmk1+/-;tmk4*, *gABP1;abp1;abl1;tmk1+/-;tmk4* and *gABL1;abp1;abl1;tmk1+/-;tmk4* lines. Scale bar, 2 mm.(D) Quantitative analysis of hypocotyl length for seedlings shown in (C). Data are mean ± SD (n=23 independent seedlings).(E) Defects in auxin-induced PC interdigitation in the *abp1;abl1;tmk1+/-;tmk4* mutant were rescued by either ABP1 (*gABP1;abp1;abl1;tmk1+/-;tmk4*) or ABL1 (*gABL1;abp1;abl1;tmk1+/-;tmk4*). Scale bar, 20 µm.(F) Quantitative analysis of PC interdigitation (lobe number/cell) shown in (E). (n>267 independent cells). ns denotes not significant; \*\*\**p < 0.001*; \*\*\*\**p* < *0.0001*; one-way ANOVA.(G-H) Auxin-induced ROP2 and ROP6 activation was abolished in the *abp1;abl1;tmk1+/-;tmk4* mutants and rescued by ABP1. Protoplasts from Col-0, *abp1;abl1;tmk1+/-;tmk4* and *gABP1;abp1;abl1;tmk1+/-;tmk4* were treated with 50 nM NAA for 10 mins. Quantitative analyses of relative ROP2 and ROP6 activity (H) are performed as described in Figure 1(E-F). Data are mean ± SD (n=3 independent experiments). ns denotes not significant; \**p < 0.05*; one-way ANOVA.

### ABL1/ABP1 and TMK1 act as co-receptors for extracellular auxin

Because of auxin-dependent physical and functional interactions between TMKs and the putative auxin binding proteins ABL1/ABL2, we hypothesize ABLs and TMKs form auxin receptor complexes on the cell surface. To test this hypothesis, we first determined whether ABL1 indeed directly binds auxin as predicted from the presence of a putative auxin binding pocket. Flag-tagged ABP1 and ABL1 fusion proteins, transiently expressed in and purified from *Arabidopsis* protoplast, were used for auxin binding assays using the microscale thermophoresis (MST) method. ABP1 proteins bound IAA with a K_*D*_ of 4.7 μM, whereas ABP1-5 proteins barely bound IAA as reported ^3,42^ (Figures 6A-C). ABL1 proteins bound IAA with a K_*D*_ of 1.7 μM, whereas the ABL1-M2 mutant protein harboring mutations in the auxin binding pocket showed no binding with IAA (Figures 6A-C). Consistent with low auxin binding of both ABP1-5 and ABL1-M2, we found *gABP1-5* and *gABL1-M2* could not complement *abp1-TD1;abl1-1;abl2-1* (*abp1;abl1/2*) mutant phenotype (Figures 3A and 3B), and *gABL1-M2* failed to complement the morphological defects in *abl1-1; abl2-1* (*abl1/2*) mutant (Figure 6D), indicating that auxin binding to ABP1 and ABL1 is essential for their physiological functions. Inactive auxin analogs such as BA (benzoic acid) and L-Trp (L-tryptophane) did not bind to either ABP1 or ABL1 (Figure S5A), confirming that ABL1 specifically binds active auxin, as shown for ABP1 ^3^.

**Figure 6.**
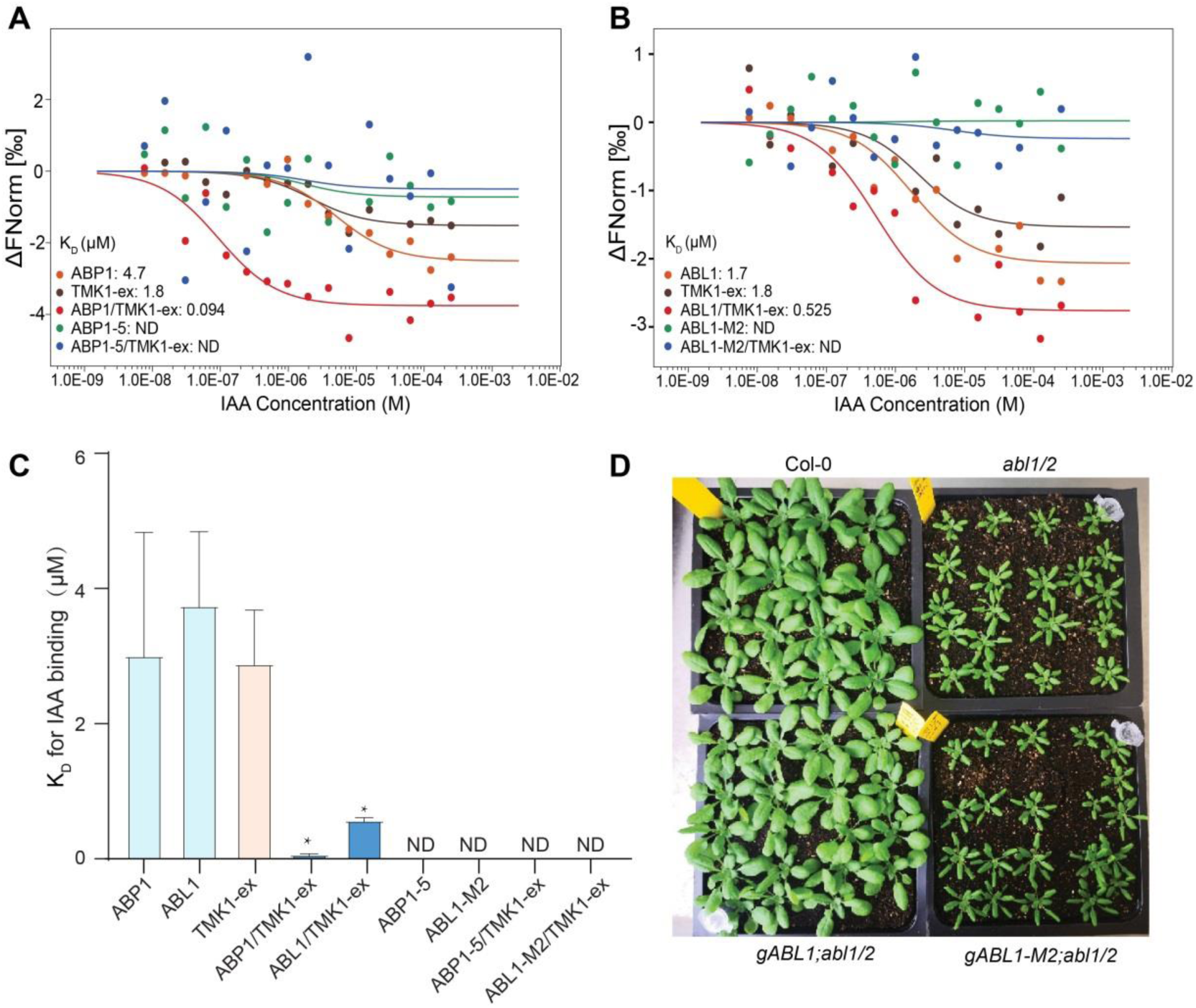
ABP1/ABL1 and TMK1 bind auxin synergistically. (A-B) Auxin (IAA) binding to ABP1, ABL1, TMK1-ex, ABP1-5, and ABL1-M2 was measured by MST (microscale thermophoresis). Data points indicate the difference in normalized fluorescence (‰) generated by no-liganded or liganded fluorescently labeled proteins, and the curves show calculated fits. Data are representatives of three independent experiments.(C) Quantification of binding affinity between IAA and fluorescently labeled ABP1, ABP1-5, ABL1, ABL1-M2, and TMK1-ex by MST in (A-B). ND denotes not detectable. Data are mean ± SD (n=3 independent experiments). \**P* ≤ 0.05; one-way ANOVA. (D) *gABL1* but not *gABL1-M2* complements the *abl1/2* mutant growth defects.

As previously reported, we found that ABP1 binds NAA with nearly 100 folds higher affinity (K_*D*_ = 35 nM) compared with IAA. Similarly, ABL1 showed a stronger binding affinity with NAA (K_*D*_ = 365 nM) than IAA. As expected, both ABP1-5 and ABL1-M2 showed dramatically reduced binding with NAA (Figure S5B), supporting direct interactions between ABL1/ABP1 and auxin. The auxin binding abilities of both ABP1 and ABL1 were further confirmed by the mammalian cell-expressed proteins. His-tagged ABP1 and ABL1, expressed in and purified from HEK293T cells, bound NAA with a K_*D*_ of 11.9 μM and 17.1 μM, respectively, whereas His-tagged ABP1-5 and ABL1-M2 failed to bind NAA (Figure S5C). Compared with *Arabidopsis* protoplast-expressed proteins, the lower auxin binding affinity for HEK293T cell-expressed ABL1 and ABP1 may result from their possible incorrect folding and/or lack of proper modifications such as glycosylation ^41,43^ in HEK293T cells, and/or from some factors co-purified from *Arabidopsis* protoplasts.

Given the auxin-dependent interaction between TMKs and ABLs/ABP1, we speculated that the extracellular domains of TMK proteins could also participate in auxin perception. We found that HA-tagged TMK1-ex fusion protein transiently expressed in and purified from *Arabidopsis* protoplasts was also able to bind IAA with a K_*D*_ of 1.8 μM and NAA with a K_*D*_ of 7.2 μM (Figures 6A and S5B), respectively, but was unable to bind inactive auxin analogs such as benzoic acid and L-tryptophane (Figure S5A). HEK293T cells-expressed His-tagged TMK1-ex bound NAA with a K_*D*_ of 71.4 μM (Figure S5C). Thus, we demonstrated that the extracellular domain of TMK1 directly binds auxin specifically.

Based on their capacity to bind auxin and to interact with each other in an auxin-dependent manner, we propose that TMK1 and ABLs/ABP1 act as co-receptors for the apoplastic auxin. Many co-receptors bind their ligands synergistically, e.g., TIR1-IAA7, PYL-PP2C, BRI1-BAK1, FLS2-BAK1^9,44–47^. Indeed the mixture of TMK1-ex with either ABP1 or ABL1, exhibited greatly higher IAA binding affinity with a K_*D*_ of 94 nM or 525 nM, respectively, compared with either ABP1/ABL1 or TMK1-ex itself (Figures 6A-C), demonstrating a synergistic auxin binding between the TMK1 and ABP1/ABL1 co-receptors, similar to the TIR1-Aux/IAA auxin coreceptor system ^9^. Altogether, our combined genetic, biochemical and cell biological data clearly indicate that ABL1/ABP1 and TMKs act as co-receptors for apoplastic auxin in plants.

## DISCUSSION

### ABL1 and ABL2 belong to a family of ancient apoplastic auxin receptors

The existence of apoplastic auxin receptors has been a Holy Grail in plant biology. For nearly half a century, the role of ABP1 as an apoplastic auxin receptor has been debated for various reasons ^7,36,48–50^, although recent studies have resurrected the function of ABP1 by demonstrating its role in auxin-mediated rapid phosphorylation and vasculature canalization in *Arabidopsis* ^3,23,51^. Because ABP1 is primarily localized to ER and its knockout mutants lack obvious morphological phenotypes, the extracellular auxin receptor remains puzzling ^23^.

Our findings here have resolved this century-long puzzle by demonstrating that ABL1 and ABL2 are new apoplastic auxin receptors. They belong to a new family of ancient auxin-binding proteins that are exclusively localized to the apoplast, and also regulate a wide range of cytoplasmic auxin responses and functionally overlap with but are distinct from ABP1 in *Arabidopsis*. These apoplastic proteins bind auxin at physiological concentrations, and their auxin-binding activity is required for their biological functions in the regulation of auxin responses and plant development. Importantly they physically and functionally act together to regulate cytoplasmic auxin responses such as ROP GTPase activation and PIN polarization. Hence, they are bonafide receptors for extracellular auxin (Figure 7).

**Figure 7.**
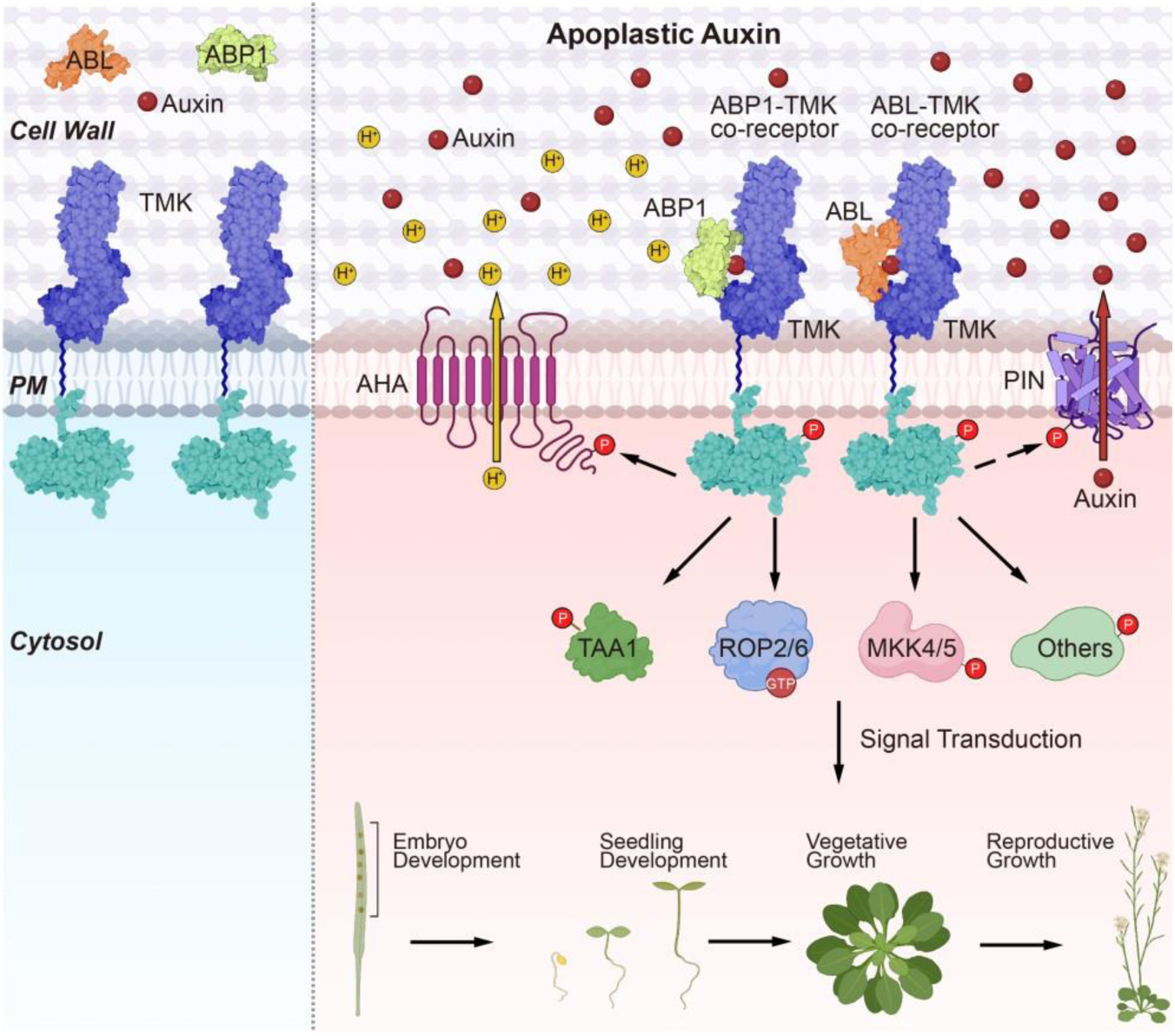
A model for the action of ABLs/ABP1 and TMKs as co-receptors for apoplastic auxin. The extracellular domain of TMK1 synergizes with both ABP1 and ABL1 in auxin binding and interacts with the apoplast-localized ABP1 and ABLs in an auxin-dependent manner. Thus, ABLs/ABP1 and TMKs form an extracellular auxin co-receptor system. Upon auxin perception, the activated TMK kinase domain directly phosphorylates a series of effectors that regulate many developmental processes from embryogenesis to reproductive development in *Arabidopsis* ^3,14,31–35^ (Figure S4 and 5).

ABL1 and ABL2 belong to a superfamily of cupin-like proteins (GLPs). Although ABP1 is also a member of this superfamily, it shares little sequence similarity to ABL1 and ABL2 except for the auxin binding pocket. However, many GLP members are much more closely related to ABL1 and ABL2 than ABP1 and are deeply conserved in plants, suggesting that at least some of GLPs may be functionally redundant with ABL1 and 2. This is consistent with our observations that both *abl1-1; abl2-1* and *abp1-TD1; abl1-1;abl2-1* mutants are conditional (Figures 3 and 4). This is in contrast to condition-independent severe developmental defects in *tmk1;tmk2;tmk3;tmk4* and *abp1;abl1;tmk1-/+;tmk4* (Figure 5) ^15^. Hence further studies are needed to identify other ABLs that interact with TMKs and function as co-receptors for extracellular auxin.

GLPs that are closely related to ABLs are conserved in bryophytes and green algae, so are TMKs ^52–54^. It is interesting that ultrafast auxin responses including auxin-induced phosphorylation that is independent of the TIR1/AFBs are also deeply conserved ^51,55^. Thus, apoplastic ABL/TMK-based perception of extracellular auxin most likely controls this ancient auxin signaling mechanism in plants.

### ABL1 and ABL2 compensate for ABP1 in the regulation of rapid auxin responses

As mentioned above, lack of morphological phenotypes in *abp1* knockout mutants and of sequence homology-based homologs of ABP1 at least in part contributed to the debates on ABP1’s role in auxin responses ^36^. How can this reconcile with the fact that ABP1 binds auxin at physiological levels, interacts with TMKs in an auxin-dependent manner, and is required for auxin-induced rapid global phosphorylation ^3,55^? We found that all three auxin-binding proteins (ABP1, ABL1, and ABL2) interact with TMK1 in an auxin-dependent manner, but among all possible multiple mutants only the *abl1-1;abl2-1* double mutant exhibits obvious morphological phenotypes and the *abp1-TD1; abl1-1;abl2-1* tripe mutant shows the most severe phenotypes. These results together with other findings here clearly demonstrate that ABL1 and ABL2 are major apoplastic auxin receptors that functionally compensate for the minor but important function of ABP1 in the perception of extracellular auxin, thus explaining the aforementioned discrepancy. Hence this work finally puts the ABP1 controversy in auxin signaling to an end.

Meanwhile, our work points out a new perspective on functional redundancy that is not simply based on protein sequence similarity, but also on protein structure similarity, providing new insight into gene compensation in plants.

### Auxin binding proteins (ABP1 and ABL1/ABL2) and TMKs are co-receptors for apoplast auxin in plants

Our findings here have finally uncovered a mechanism for the perception of apoplastic auxin in plants (Figure 7). Previous work shows that TMKs are essential for auxin responses originating at the plasma membrane that impact a wide range of developmental processes from embryogenesis to organ formation and growth ^14,33,37,56,57^, but how TMKs perceive auxin is unclear. We demonstrate that the extracellular domain of TMK1 interacts with the apoplast-localized ABL1 and ABL2 in an auxin-dependent manner. More importantly, the extracellular domain of TMK1 itself binds auxin and synergizes with both ABP1 and ABL1 in auxin binding. Together with the genetic interaction between ABP1/ABLs and TMKs, these results indicate that ABLs/ABP1 and TMKs are co-receptors for apoplastic auxin, establishing an extracellular auxin sensing system (Figure 7). This system is most likely deeply conserved, as homologs for both TMKs and ABP1/ABLs exist in land plants and green algae ^52–54,58^ and is widely used to control various important auxin responses given the functions of TMKs in auxin responses (Figure 7) ^14,15,33,35^. Furthermore, auxin rapidly induces the phosphorylation of a large set of proteins that is commonly ABP1- and TMK-dependent, including the deeply conserved RAF-like protein ^3,51,55^. The diverse array of TMK targets and the various functions of downstream pathways suggest the importance of monitoring the dynamic of apoplastic auxin and various levels of extracellular auxin. Future studies should elucidate at the structural level how auxin modulates the ABLs/ABP1-TMK interaction and is perceived by this apoplastic co-receptor system. Since apoplastic auxin dynamics need to be tightly linked to the fluctuation of intracellular auxin levels, it will also be important to understand how the perception and signaling of apoplastic auxin are coordinated with intracellular auxin signaling pathways controlled by the TIR1/AFB auxin receptors.

## ACKNOWLEDGEMENTS

This work was mainly supported by the National Natural Science Foundation of China (Grant 32130010, 31422008), startup funds from both FAFU (Fujian Agriculture and Forestry University) and PSC (Shanghai Center for Plant Stress Biology, and Center for Excellence in Molecular Plant Sciences, Chinese Academy of Sciences) to T. X., and in part supported by a grant from US National Institute of General Medical Sciences (GM100130) to ZY. Figure 7 was drawn with BioRender.com.

## AUTHOR CONTRIBUTIONS

T. X., Z. Y., and Y. Y. initiated the project and designed the experiments. Y. Y., W. T., W. Lin. performed the majority of experiments. W. Li. and X. Z. performed plasmid constructions, phenotype analysis, immuno-staining, FRET analysis, and AlphaFold analysis. Y. L., R. C., R. H., J. M., R. Z., helped with the preparation of the mutant materials. P. P. helped with pavement cell analysis. J. L., performed transcriptome analysis, Q. G., helped with the IP-MS, W. C and L. J performed immunogold staining. T. X. and Z. Y. wrote the manuscript.

## MATERIALS AND METHODS

### Plant materials and growth conditions

Col-0 was used as an *Arabidopsis* wild-type ecotype, and all mutants and transgenic lines were produced in this background. The *abp1-TD1*, *tmk1-1, tmk4-1, tmk1-1;tmk4-1*, *BRI1-GFP*, *TMK1-Flag*(*pTMK1::TMK1-Flag*), and *TMK1-GFP* (*pTMK1::TMK1-GFP*) lines have previously been described ^15,33,34,36^. The T-DNA insertion line *abl1-1* (GABI_820G11) was obtained from the European *Arabidopsis* Stock Centre (NASC, Nottingham, UK). The *abl2-1* single mutant was generated using the CRISPR-Cas9 technology ^59^. We designed two single-guide RNA (sgRNA) sequences targeting ABL2 to increase editing efficiency. The CRISPR constructs were transformed into wild-type (Col-0) by floral dipping using *Agrobacterium tumefaciens* GV3101^60^. The Cas9-free homozygous mutants were obtained by GFP fluorescent screening and were confirmed by direct sequencing of PCR products from the offspring in the T3 generations. Double and higher-order mutants of *abp1-TD1*, *abl1-1*, *abl2-1*, *tmk1-1,* and *tmk4-1* were generated by standard genetic crossing and appropriate genotypes were selected using gene-specific mutant genotyping primers. The ABP1-5(H94Y) and ABL1-M2(H100AH102A) point mutations were obtained by using PCR-based site-specific mutagenesis ^61^. To generate the transgenic *pABP1::ABP1;abp1-TD1*, *pABP1::ABP1-5(H94Y);abp1-TD1*, *pABP1::ABP1;abp1-TD1;abl1-1;abl2-1* and *pABP1::ABP1-5(H94Y);abp1-TD1;abl1-1;abl2-1* lines, a 5.2-kb genomic fragment containing the promoter region, the coding region, and the 3’ untranslated region was amplified by PCR and inserted into a pDONR-Zeo vector, subsequently transferred into a pGWB401 destination vector and then transformed into the *abp1-TD1 and abp1-TD1;abl1-1;abl2-1* mutants, respectively. To generate the transgenic *pABL1::ABL1;abp1-TD1;abl1-1, pABL1::ABL1;abl1-1;abl2-1*, *pABL1::ABL1-M2* (*H100AH102A*)*;abl1-1;abl2-1, pABL1::ABL1;abp1-TD1;abl1-1;abl2-1* and *pABL1::ABL1-M2* (*H100AH102A*)*;abp1-TD1;abl1-1;abl2-1* lines, a 4.1-kb genomic fragment containing the promoter region, the coding region and the 3’ untranslated region was amplified by PCR and inserted into a pDONR-Zeo vector, subsequently transferred into a pGWB401 destination vector and then transformed into the *abp1-TD1;abl1-1*, *abl1-1;abl2-1* and *abp1-TD1;abl1-1;abl2-1* mutants, respectively. The promoter region of the pDONR-Zeo vector containing *pABP1::ABP1* was replaced with *ABL1* promoter, transferred into a pGWB401 destination vector and then transformed into the *abp1-TD1;abl1-1;abl2-1* mutant to generate the transgenic *pABL1::ABP1;abp1-TD1;abl1-1;abl2-1* lines, *pABP1::ABP1;abp1-TD1* and *pABL1:ABL1;abp1-TD1;abl1-1* was used to cross with *tmk1-1;tmk4-1* to generate *pABP1:ABP1;abp1-TD1;abl-1;1tmk1-1-/+tmk4-1* and *pABL1:ABL1;abp1-TD1;abl1-1;tmk1-1-/+tmk4-1*, respectively. All primers used for transgenic line generation are listed in Supplementary Table 1.

Most of the plant materials were grown in a plant growth room or chamber (PERVICAL AR-66L3) at 23°C, 70% humidity, and 75 μmol m^-2^ s^-1^ light intensity under a 16 h light/8 h dark photoperiod in Fuzhou, Fujian, China, unless otherwise indicated. Mutants and complementary lines related to ABL1 and ABL2 that were used for phenotype analysis (Figure 3, Figure 4, and Figure 6D) were planted in a growth room at 21°C, 65% humidity, and 75 μmol m^-2^ s^-1^ light intensity under a 16 h light/8 h dark photoperiod in Nanjing, Jiangsu, China. These growth phenotypes were reproduced at University of California, Riverside, CA, USA; Shanghai Center for Plant Stress Biology, CAS, Shanghai, China. The phenotypes were mild or not obvious at Fujian Agriculture and Forestry University, Fuzhou, China.

For *Arabidopsis* seedling growth on agar media, seeds were surface sterilized with 75% (v/v) ethanol (containing 0.05% Triton X-100) for 8 mins, rinsed with sterile water three times, and then placed on the plates with 1/2 MS medium containing 0.5% sucrose (w/v) and 0.8% (w/v) agar at pH 5.7, stratified for 48 hours before moved to growth chambers.

### Auxin treatment of cotyledons to promote PC interdigitation

After two days of vernalization in 1.5 ml sterile water, the surface-sterilized seeds were evenly placed into 6-well plates (less than 30 seeds per well) containing 2 ml semi-liquid 1/2 MS (1/2 MS with 0.2% agar) with or without 20 nM NAA treatment. The 6-well plates were placed in an incubator and grown for 3 days before imaging.

### Analysis of *Arabidopsis* cotyledon pavement cell shape

*Arabidopsis* seedlings were stained with propidium iodide (PI), and pavement cells in the same region that just below the tip of cotyledons were imaged and photographed on a Leica SP8 confocal microscope. The morphological analysis of pavement cells was performed by PaCeQuant ^62^. The related parameters are as follows: phases to run, segmentation_and_features; operation mode, Batch; Pixel calibration mode, Auto; border contrast, bright_on_dark; heuristic for Gap closing, Watershed; Minimal size of cells, 2500, unit for size thresholds, PIXELS; Maximal size of Cells, 1000000. Cell area, neck width, lobe length, lobe number, and margin roughness were outputs for further analysis.

### ROP activity measurement

Western blotting-based ROP activity measurements were performed as described ^63^. Total ROP and active ROP proteins that were pulled down by MBP-RIC1 (GTP-bound ROP) were detected by western blotting analysis using anti-ROP2 (ABclonal), anti-ROP6 (ABclonal), and horseradish peroxidase-conjugated rabbit antibodies. For ROP activity assays in protoplasts, protoplasts were isolated from 3-week-old leaves. Isolated protoplasts were treated with 20 nM NAA and frozen in liquid nitrogen. For ROP activity assays in 3-week-old leaves, 0.4-0.5 g samples were collected for ROP activity measurements. Image J was used to quantify the amounts of ROPs.

### Auxin-induced rapid hypocotyl elongation

For analyzing auxin-induced hypocotyl elongation in seedlings, 5-day-old seedlings grown in 1/2 MS medium were transferred into 1/2 MS medium containing indicated concentrations of NAA and incubated for additional 48 hours at 23°C, and 75 μmol m^-2^ s^-1^ light intensity under a 16 h light/8 h dark photoperiod. Hypocotyl lengths were measured by Image J after treatment.

### Subcellular localization of ABL1 by immuno-gold labeling and transmission electron microscopy (TEM)

For TEM analysis, sample preparation, high-pressure freezing, freeze substitution, resin embedding, and ultramicrotomy were performed as described previously ^64^. 5-day-old wild-type *Arabidopsis* roots or 3-week-old true leaves were frozen in a high-pressure freezer (EM ICE, Leica), followed by freeze substitution in dry acetone containing 0.4% uranyl acetate at −85°C in a freeze-substitution unit (EM AFS2, Leica). After three acetone rinses at −85°C, the samples were removed from the planchets and slowly infiltrated stepwise with increasing concentrations of HM20 resin over 96 hours before ultraviolet polymerized at −35°C. Ultra-thin sectioning was performed in an ultramicrotome unit (EM UC7, Leica).

Immuno-gold labeling was performed as previously described ^64,65^ with anti-ABL1 antibodies diluted at 1: 25 in 1% BSA, and 10 nm gold-coupled rabbit secondary antibodies at 1: 40 dilution in 1% BSA, followed by a post-staining procedure using aqueous uranyl acetate/lead citrate. Samples were observed under Hitachi H-7650 TEM (Hitachi High-Technologies Corporation, Japan) with a charge-coupled device camera operating at 80 kV.

### Co-immunoprecipitation analysis

The transgenic TMK1-GFP line was used to analyze interactions between TMK1 and ABL1 or ABL2. A BRI1-GFP line was used as a negative control. After vernalization, seeds were cultured in liquid 1/2 MS on a 6-well plate (less than 50 seeds per well) for 7 days. The seedlings were washed with 2 ml new liquid 1/2 MS for 1 h, and then treated with 2 ml 1/2 MS containing different concentrations of NAA. After 15 min of treatment, *Arabidopsis* seedlings were dried on filter paper. A total of 0.2 g of seedlings for each sample was collected and frozen in liquid nitrogen. The same procedure was performed for time course analysis. *Arabidopsis* seedlings were treated with 200 nM NAA for 0 min, 5 mins, 15 mins, and 30 mins, respectively. Then 0.2 g samples were collected and frozen in liquid nitrogen.

The samples were ground to powder in liquid nitrogen, and incubated in 500 µL extraction buffer (10 mM HEPES at pH 7.5, 100 mM NaCl, 1 mM EDTA, 10% (vol/vol) glycerol, 0.5% Triton X-100 (v/v), and protease inhibitor mixture from Roche). After lysis for 15 mins on ice, the samples were centrifuged at 12,470 g for 15 mins at 4°C. 40 µL supernatant was collected for input analysis, and the remaining was supplied with 740 µL extraction buffer and then incubated with GFP-Trap (Chromotek, 10 µL for each sample) for 2 hours with gentle shaking. The beads were collected and washed three times with a washing buffer (10 mM HEPES at pH 7.5, 100 mM NaCl, 1 mM EDTA, 10% glycerol, v/v, and 0.1% Triton X-100, v/v) and once with 50 mM Tris-HCl at pH 7.5. The immunoprecipitated proteins were analyzed by western blotting analysis with anti-ABL1 and anti-ABL2 antibodies (1:2000 dilution, 4°C, overnight).

Polyclonal *Arabidopsis* ABL1 and ABL2 antibodies were generated by ABclonal Biotechnology (China, Wuhan). In brief, synthetic peptides for ABL1 (24-38aa, VANLKRAETPAGYPC) and ABL2 (32-49aa,GPQSPSGYSCKNPDQVTEC) coupled to keyhole limpet hemocyanin (KLH) were injected into rabbits as antigens, respectively. Antibodies were affinity-purified from immunized rabbit serum using a Sulfo-Link matrix (Pierce). The specificity of the ABL1 and ABL2 antibodies were tested by western blotting analysis (1:2000 dilution for both antibodies) of protein extracts from the *abl1-1* and *abl2-1* mutants, respectively.

### FRET analysis of the interaction between TMK1-ex and ABL1

*Agrobacterium tumefaciens* GV3101 carrying *35S::mCherry-TMK1-ex* or *35S::ABL1/ABL1-M2-GFP* plasmid were harvested from overnight culture and resuspended with buffer (10 mM MgCl2, 10 mM MES, 200 mM Acetosyringone, pH 5.7) at OD600 = 0.8 for tobacco leaf infiltration. Two days after infection, the tobacco leaves were dipped into 100 nM NAA or mock solution for 5 mins before imaging. The FRET efficiency was analyzed by FRET Sensitized Emission methods ^66^. In brief, the ABL1-GFP/ABL1-M2-GFP only, mCherry-TMK1-ex only sample, and the FRET samples were imaged with the same microscopic setting on Leica SP8 confocal laser scanning microscope. Donor and FRET channels were excited with 70% white light laser at 488 nm (10%), and the emissions were collected by HyD detector using 499-551 nm and 600-650 nm respectively. Acceptor channel was excited with 70% white light laser at 561 nm (50%), and the emissions were collected by HyD detector using 600-650 nm). A segmented line was drawn along the cell boundary region of the tobacco pavement cell to measure the mean signal intensity for each channel with Image-Pro Plus (http://www.mediacy.com/imageproplus) and LAS-X (Leica). The correction factors β, α, γ and δ were calculated with the Donor and Acceptor only reference samples, and the FRET-SE efficiency was calculated with the equation described by van Rheenen ^66^.

To generate the FRET-SE efficiency heatmap image in Figure 2G, the cell boundary region of tobacco pavement cell (dotted lines) was cropped as the region of interest from Donor, FRET, and Acceptor channels. Then these ROIs were processed by the image calculator module of ImageJ with the *E* _(FRET-SE)_ equation described by van Rheenen ^66^.

### ABL1 and ABP1 auxin binding assays

ABL1 and ABP1 fusion proteins for auxin binding assay were expressed in and purified from both *Arabidopsis* protoplasts and mammalian cells. Protoplasts were prepared according to the protocol described by Im and Yoo ^67^. Maxiprep DNA for transient expression was prepared using the Invitrogen PureLink Plasmid Maxiprep Kit. 2x10^5^ protoplasts were transfected with HBT vector harboring ABP1 and ABP1-5 with Flag tag insertion after the glycine 120 ^68^, TMK1-ex with HA tag and ABL1, ABL1-M2 with Flag tag at the C-terminal, and incubated at room temperature for 10 hours. The protoplasts were collected and stored at −80°C for future usage.

For protein affinity purification from protoplasts, 50 ml of protoplasts containing expressed fusion proteins were collected and lysed with extraction buffer (50 mM Tris-HCl pH 7.4, 150 mM NaCl, 5 mM EDTA, 0.5% Triton X-100 with protease inhibitor and phosphatase inhibitor) on ice. The extracts were centrifugated at 12000 g for 10 mins, and the supernatants were incubated with HA agarose beads (Sigma-Aldrich, A2095) or Flag-agarose beads (Sigma-Aldrich, A2220) at 4°C for 3 hours to immunoprecipitate TMK1-ex-HA, ABP1-Flag, or ABL1-Flag proteins. The agarose beads were washed and resuspended with extraction buffer with 0.1% Triton X-100 3 times, and then washed with 50 mM Tris-HCl buffer (pH 7.8). One 10th of the beads were used for immunoblot analysis with anti-HA and anti-Flag antibodies, respectively. The remaining agarose beads were eluted with IgG-Eluted buffer. The eluted proteins were used for MST analysis (described below).

For protein purification from mammalian cells, coding sequences for ABP1, ABP1-5, ABL1, and ABL1-M2 were synthesized by GENEWIZ. To facilitate protein expression in mammalian cells, codon optimization was done while keeping the amino acid sequence unchanged. Synthesized sequences were cloned into modified pcDNA3.4 vectors, containing an N-terminal secretion signal peptide and a C-terminal 6×His-tag. All proteins were expressed using the HEK293T expression system. HEK293T cells were routinely cultured as previously described ^69^. Transfections were performed using the PEI-max ^70,71^ method. The supernatant was collected via centrifugation on the 7th day after infection. The supernatant flowed through Ni-NTA (Novagen). Bound proteins were eluted in a buffer containing 25 mM Tris-HCl pH 8.0, 150 mM NaCl, and 250 mM imidazole and further purified by size exclusion chromatography (Hiload 16/60 Superdex 200 prep grade, GE Healthcare) in a buffer containing 10 mM Bis-Tris pH 6.0, 100 mM NaCl. All proteins were diluted to a final concentration of 10 μM in a buffer containing 10 mM Bis-Tris pH 6.0, and 100 mM NaCl ^69^.

The binding affinity between auxin, BA (benzoic acid) and L-Trp (L-tryptophane), and the ABP1, ABP1-5, ABL1, ABL1-M2, TMK1-ex, and FLS2-ex proteins was measured by the microscale thermophoresis (MST) method ^72^. 10 μM purified wild-type and mutated ABP1 and ABL1, TMK1-ex and FLS2-ex proteins were labeled with a fluorescent dye using a MO-L001 MonolithTM Protein Labelling Kit RED-NHS (Amine Reactive). Auxin in a range of concentrations (5 mM to 0.153 μM NAA for mammalian cell expressed proteins, 250 μM to 0.00763 μM BA, 250 μM to 0.00763 μM L-Trp, 250 μM to 0.00763 μM IAA and 50 μM to 0.00153 μM NAA for *Arabidopsis* protoplast expressed proteins) was incubated with labeled proteins (60 nM) in buffer containing (20 mM Mes-NaOH pH 5.7, 150 mM NaCl, 10 mM ZnCl2 and 0.001% Tween 20) for 30 mins at 4°C. Approximately 4-6 μl of each sample was loaded into silica capillaries (Monolith™ NT.115 Standard Treated Capillaries, MO-K002) and microthermophoresis was carried out in a Monolith NT.115 instrument using 20% LED power and 40% MST. Measurements were performed repeatedly on independent protein preparations to ensure reproducibility. The KD values were calculated using the mass action equation via the NanoTemper MO. Affinity Analysis software (NanoTemper Technologies).

### Quantification and statistical analysis

All data were analyzed using the GraphPad Prism9 software. Significant differences were determined using Student’s *t*-test, one-way ANOVA, and two-way ANOVA. Multiple comparisons were determined by Tukey–Kramer HSD (*p* < 0.05).

## FIGURE LEGENDS

**Figure S1. *gABP1-5;abp1* lines show defects in root growth and gravitropic response, cotyledon development, and leaf morphology** (A) Western blotting analysis of ABP1 protein levels in Col-0, *abp1*, *gABP1-5;abp1* and *gABP1;abp1* lines. (B) PIN1 localization in 5-day-old seedling roots of Col-0, *abp1* and *gABP1-5;abp1* lines. Scale bar, 10 µm.(C) Quantification of the PIN1 polarity shown in (B) by the fluorescence ratio of apical and basal (AB)/lateral (L) membranes sides of PIN1. Data are mean ± SE (n≥ 40 cells from more than 6 roots for each line). ns denotes not significant; \**p* < *0.05*; one-way ANOVA. (D) Root phenotype from 7-day-old seedlings of Col-0, *abp1*, *gABP1-5;abp1*, *gABP1;abp1* lines. Scale bar, 0.5 cm.(E) Quantification of the root length of the seedlings in (D). n=26 independent seedlings. ns denotes not significant; \*\*\*\**p < 0.0001*; one-way ANOVA.(F) Gravitropic responses of Col-0, *abp1*, *gABP1-5;abp1* and *gABP1;abp1* lines. Scale bar, 0.2 cm.(G) Quantification of root bending angles in Col-0, *abp1*, *gABP1-5;abp1* and *gABP1;abp1* lines in (D). n=32 independent seedlings. ns denotes not significant; \*\*\*\**p* < *0.0001*; one-way ANOVA.(H) Abnormal cotyledon ratios of Col-0, *abp1*, *gABP1-5;abp1* and *gABP1;abp1* lines. Scale bar, 0.2 cm.(I) The morphologies of 3-week-old soil-grown Col-0, *abp1*, *gABP1-5;abp1* and *gABP1;abp1* lines. Scale bar, 1 cm.

**Figure S2. ABL1 has a similar structure to ABP1 and apoplastic localization,** (A) Sequence alignment of auxin binding pockets of ABP1, ABL1, and ABL2.(B) Binding mode of ABP1, ABL1, and ABL2 to NAA by molecular docking. The white and purple dashed lines indicate the hydrogen bonds and the π-π stacking, respectively.(C) Validation of ABL1 and ABL2 antibodies in Col-0, *abl1*and *abl2* lines by western blotting analysis.(D) ABL1 is localized to the apoplast in old leaves and roots by the immune colloidal gold technique. CE, cell wall and extracellular region. ECE, regions except for cell wall and extracellular region. Scale bar, 500 nm. (E-F) Quantification of colloidal gold particles in old leaves (E) and root (F). CE, cell wall and extracellular region. ECE, regions except for cell wall and extracellular region. Data are mean ± SD (n>10 regions). ns denotes not significant; \*\*\*\**p* < *0.0001*; one-way ANOVA.(G) ABL1 localization was determined by immuno-staining using primary anti-ABL1 antibody and secondary Dylight488 Goat anti-Rabbit IgG antibody. Scale bar, 10 µm.

**Figure S3. The generation and characterization of *abl1* and *abl2* mutant** (A) Schematic diagrams of ABL1 T-DNA insertion (*abl1-1*) and ABL2 CRISPR target points. Black and white boxes represent exons and untranslated regions, respectively. * indicates the early stop codon in the *abl2-1* mutant.(B) Validation of *abl1-1* mutant by RT-PCR.(C) PIN1 localization in 5-day-old seedling roots of Col-0 and *abp1;abl1/2* lines. Scale bar, 20 µm.(D) Quantification of the PIN1 polarity shown in (C) by the fluorescence ratio of apical and basal (AB)/lateral (L) membranes sides of PIN1. Data are mean ± SE (n≥ 48 cells from more than 6 roots for each line). \*\**p* < *0.01*; Student *t-*test.(E-F) Expression atlas of ABP1, ABL1, ABL2, TMK1 and TMK4 in different tissues. TPM (transcriptomic) and iBAQ (proteomic) data obtained from Mergner et al., ^73^ were Z-score centered and shown by heatmap.

**Figure S4. Defects of inflorescence phenotype and root development in *abp1;abl1tmk1+/-tmk4*** (A) The morphological phenotype of 6-week-old soil-grown Col-0, *abp1;abl1*, *tmk1+/-;tmk4*, *abp1;tmk1+/-;tmk4*, *abl1;tmk1+/-;tmk4*, *abp1;abl1;tmk1+/-;tmk4*, *gABP1;abp1;abl1;tmk1+/-;tmk4* and *gABL1;abp1;abl1;tmk1+/-;tmk4* lines. Scale bar, 5 cm. (B) Silique phenotype of 6-week-old soil-grown Col-0, *abp1;abl1*, *tmk1+/-;tmk4, abp1;tmk1+/-;tmk4*, *abl1;tmk1+/-;tmk4*, *abp1;abl1;tmk1+/-;tmk4*, *gABP1;abp1;abl1;tmk1+/-;tmk4* and *gABL1;abp1;abl1;tmk1+/-;tmk4* lines. Scale bar, 1 cm.(C) Quantification of PC interdigitation depicted by margin roughness is shown in Figure 5 (E). n>267 independent cells for each treatment. ns denotes not significant; \*\*\*\**p* < *0.0001*; one-way ANOVA. (D) PIN1 localization in 5-day-old seedling roots of Col-0, *abp1;abl1* and *abp1;abl1;tmk1+/-;tmk4* lines. Scale bar, 10 µm.(E) Quantification of the PIN1 polarity shown in (D) by the fluorescence ratio of apical and basal (AB)/lateral (L) membranes sides of PIN1. Data are mean ± SE (n≥ 44 cells from more than 6 roots for each line). ns denotes not significant; \**p* < *0.05*; one-way ANOVA.(F) Root phenotype from 7-day-old seedlings of Col-0, *abp1;abl1*, *tmk1+/-;tmk4*, *abp1;tmk1+/-;tmk4*, *abl1;tmk1+/-;tmk4*, *abp1;abl1;tmk1+/-;tmk4*, *gABP1;abp1;abl1;tmk1+/-;tmk4* and *gABL1;abp1;abl1;tmk1+/-;tmk4* lines. Scale bar, 0.5 cm.(G) Quantification of the root length of the seedlings in (F). n>41 independent seedlings. Different letters indicate values with statistically significant differences (*p < 0.05*; Tukey HSD).

**Figure S5. IAA and NAA binding to ABP1, ABL1 and TMK1-ex, but not ABP1-5, or ABL1-M2 measured by MST** (A-C) Data point indicated the difference in normalized fluorescence (‰) generated by no-liganded or liganded fluorescently labeled indicated proteins, and curved indicated the calculated fits. The result is representative of three independent experiments. ABP1, ABP1-5, ABL1, ABL1-M2, TMK1-ex and FLS2-ex proteins were expressed in *Arabidopsis* protoplasts (A-B) and in mammalian cells (C). (A) Quantification of binding affinity between IAA, BA, L-Trp and fluorescently labeled ABP1, ABP1-5, ABL1, ABL1-M2, TMK1-ex and FLS2-ex by MST in (A). ND denotes not detectable. Data are mean ± SD (n=3 independent experiments).(B) Quantification of binding affinity between NAA and fluorescently labeled ABP1, ABP1-5, ABL1, ABL1-M2, TMK1-ex and FLS2-ex by MST in (B). ND denotes not detectable. Data are mean ± SD (n=3 independent experiments). (C) Quantification of binding affinity between NAA and fluorescently labeled ABP1, ABP1-5, ABL1, ABL1-M2, TMK1-ex by MST in (C). ND denotes not detectable. Data are mean ± SD (n=3 independent experiments).

